# panRGP: a pangenome-based method to predict genomic islands and explore their diversity

**DOI:** 10.1101/2020.03.26.007484

**Authors:** Adelme Bazin, Guillaume Gautreau, Claudine Médigue, David Vallenet, Alexandra Calteau

## Abstract

**Motivation:** Horizontal gene transfer (HGT) is a major source of variability in prokaryotic genomes. Regions of Genome Plasticity (RGPs) are clusters of genes located in highly variable genomic regions. Most of them arise from HGT and correspond to Genomic Islands (GIs). The study of those regions at the species level has become increasingly difficult with the data deluge of genomes. To date no methods are available to identify GIs using hundreds of genomes to explore their diversity.

**Results:** We present here the panRGP method that predicts RGPs using pangenome graphs made of all available genomes for a given species. It allows the study of thousands of genomes in order to access the diversity of RGPs and to predict spots of insertions. It gave the best predictions when benchmarked along other GI detection tools against a reference dataset. In addition, we illustrated its use on Metagenome Assembled Genomes (MAGs) by redefining the borders of the *leuX* tRNA hotspot, a well studied spot of insertion in *Escherichia coli*. panRPG is a scalable and reliable tool to predict GIs and spots making it an ideal approach for large comparative studies.

**Availability:** The methods presented in the current work are available through the following software: https://github.com/labgem/PPanGGOLiN. Detailed results and scripts to compute the benchmark metrics are available at https://github.com/axbazin/panrgp_supdata.

**Supplementary information:** None.

## 1 Introduction

Horizontal gene transfer (HGT) is a major mechanism that shapes gene repertoires of bacterial species providing and maintaining diversity at the population level (Ochman *et al.*, 2000; Niehus *et al.*, 2015). This pervasive evolutionary process spreads genes between, potentially very distant, bacterial lineages (Thomas and Nielsen, 2005) and is a significant source of gene novelty (Treangen and Rocha, 2011). In prokaryotes, HGT is promoted by three main mechanisms: conjugation, transformation or transduction. Clusters of consecutive genes likely acquired by HGT are commonly described as Genomic Islands (GIs) (Hacker and Kaper, 2000). GIs are part of the flexible gene pool of bacteria and may bring an evolutionary advantage allowing adaptations to new environments or bringing new pathogenicity capacities for instance (Hacker and Carniel, 2001). They are widely distributed in pathogenic and environmental microorganisms outlining the interest that researchers have in studying their evolution and functional impact on bacterial populations.

GIs are characterized by their large size (>10 kb), a usually different G+C content compared with the rest of the chromosome for recent acquisitions (Lawrence and Ochman, 1997), and are often associated with mobile elements such as transposons, integrons, Integrative Conjugative Elements (ICEs) and prophages. Many GIs insertion sites are associated with tRNA-encoding genes and are flanked with repeat structures (Dobrindt *et al.*, 2004). Some of those insertion sites, called hotspots, are more active than the rest of the genome in terms of acquisition rate of new elements and tend to have a much more diverse gene content even between closely related individuals (Oliveira *et al.*, 2017). Since GIs carry so many genes of interest, countless methods have been published to detect and analyze them (Langille *et al.*, 2008; Lu and Leong, 2016; Bertelli *et al.*, 2018). These methods are often grouped into two categories: composition-based methods and comparative genomic-based methods. A recent review describes many of the methods that were developed in this field (Bertelli *et al.*, 2018) and benchmarks them using a curated dataset (Langille *et al.*, 2008).

Nowadays, a deluge of microbial genomes is available in public databanks with more than half a million prokaryotic genomes in Genbank (last accessed 2nd March 2020) (Sayers *et al.*, 2019). In parallel, environmental data made of Metagenome Assembled Genomes (MAGs) or Single-cell Assembled Genomes (SAGs) are increasing dramatically. Hence, conducting comparative genomics studies on hundreds to thousands of genomes has become a challenge and can lead to millions of pairwise comparisons requiring intensive computations for analysis. Accurately identifying GIs in all of those genomes that may be incomplete and fragmented is becoming crucial to get a global overview of the diversity within species.

To tackle this challenge, methods based on pangenomes could be the answer. The concept of pangenome corresponds to the entire gene repertoire of a taxonomic group (Tettelin *et al.*, 2005). A pangenome can be described by two components: the core genome, which contains genes shared by all individuals, and the accessory genome, which gathers every other genes. Lately, multiple methods have been developed to study pangenome structures and to perform comparative studies on hundreds of genomes (Fouts *et al.*, 2012; Page *et al.*, 2015; Snipen and Liland, 2015; Gautreau *et al.*, 2020). Among them, the PPanGGOLiN method proposes a representation of all genes of all genomes in a pangenome graph where nodes represent gene families and edges represent genomic neighborhood (Gautreau *et al.*, 2020). The pangenome graph is divided using a statistical model in three partitions that represent the occurrence of the gene families as following : (i) the *persistent* genome which corresponds to genes that are present in most individuals of the studied clade, (ii) the *shell* genome which groups genes that are conserved between some individuals of the group but not most and (iii) the *cloud* genome which corresponds to genes that are rare within the population and found only in one or a few individuals. The *shell* and *cloud* genomes are partitions of the accessory genome. The *persistent* genome is conceptually similar to the core genome but it is more adapted to large-scale genomic comparisons as it allows for missing genes due to ponctual evolutionary loss events or technical reasons such as assembly or gene calling artifacts (Gautreau *et al.*, 2020).

As most newly acquired genes are expected to arise from HGT events (Treangen and Rocha, 2011), it is expected that most of the genes included in the *shell* and *cloud* genomes have a non-vertical origin and are either part of GIs or plasmids. Here, we use the concept of Regions of Genome Plasticity (RGP) to refer to regions composed of *shell* and *cloud* genomes (Mathee *et al.*, 2008; Vallenet *et al.*, 2009; Ogier *et al.*, 2010). We expect RGPs to mostly consist of GIs or plasmids. In the case of significant genome reduction in the studied species, regions that have been lost in some individuals might be included in the *shell* genome and thus considered as RGPs. While GIs have previously been studied in the scope of pangenomes and were shown to include most of the variable genome in different species (Kittichotirat *et al.*, 2011; Zhu *et al.*, 2019), so far no method uses the concept of pangenome to predict them.

To study the evolution of GIs in a population, it may be of interest to look at spots of insertion within a pangenome. The name spot of insertion has been used previously to describe the variable genome of multiple individuals that was located in-between the same core genes (Lescat *et al.*, 2009; Oliveira *et al.*, 2017). A related concept can be found in the literature in the name of flexible genomic regions (fGR) which is a group of flexible genomic islands (fGI) (Chan *et al.*, 2015), a term originally used to describe the variable genome of different individuals which was located in between the same core genes and involved in similar functions (Rodriguez-Valera and Ussery, 2012). We will use the term spot throughout this paper and we do not assume that genes in a same spot have related functions.

In the present paper, we propose a new method called panRGP which detects RGPs and gathers them into spots of insertion to study the dynamics of GIs. It is a comparative genomic-based method that uses a pangenome graph reconstructed from hundreds to thousands of genomes of the same species. We benchmarked panRGP along a selection of other tools against a previously published dataset on GIs (Langille *et al.*, 2008). Finally, we illustrated its use on incomplete and fragmented genomes such as MAGs in the context of the analysis of an insertion spot in *Escherichia coli*.

## 2 Materials and Methods

### 2.1 panRGP method

The panRGP method predicts RGPs from a query genome that is annotated with a set of protein-coding genes. It uses as input a partitioned pangenome graph that is built from the genomes of related organisms usually from the same species. This graph is based on the PPanGGOLiN data structure (Gautreau *et al.*, 2020) where nodes are homologous gene families and edges indicate a relation of genetic contiguity. The different steps for the detection of RGPs are shown in Figure 1 and are detailed in subsections 2.1.1 and 2.1.2. Firstly, each gene of the query genome is assigned to the pangenome partition of its gene family and thus classified as *persistent*, *shell* or *cloud*. Secondly, a score is computed for each gene sequentially along the genome. It is based on both the gene partition and the score of the previous gene. Finally, RGPs are detected using the gene scores and correspond to sequences of *shell* and *cloud* genes possibly interrupted by few *persistent* genes. In addition, RGPs from different genomes can be grouped in spots of insertion based on their conserved flanking *persistent* genes using the pangenome graph as explained in subsection 2.1.3.

**Fig. 1.**
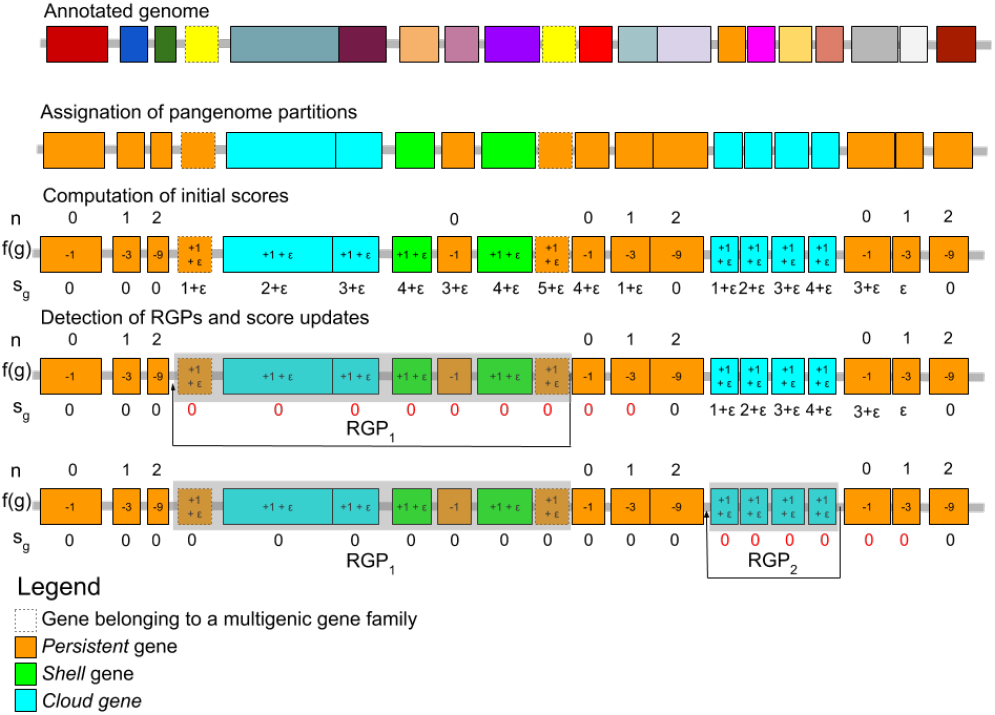
Overview of the RGP detection method. The different steps of the RGP detection method are presented. Boxes represent protein-coding genes and their color is indicative of their gene families for the annotated genome and their pangenome partition for the other genome representations. Dashed boxes indicate genes belonging to multigenic gene families. In this example, two RGP are detected. *RGP*_1_ has a score of 5, and *RGP*_2_ a score of 4. *n* values indicate for each gene the number of upstream consecutive genes classified as persistent. *f* (*g*) values indicate the result of a function that is used to compute the gene score *s_g_*. In this example, the default values for *p* and *v* parameters are used and are 3 and 1, respectively.

#### 2.1.1 Computation of initial gene scores

The initial step consists of assigning a score to each gene *g,* ∀*g* ∈ *C* where *C* is an ordered set of genes that are present on each contig of a genome assembly. The gene scores *s_g_* are computed sequentially along the contigs as follows:

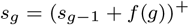

*s_g_*_−1_ is the score of previous gene on the contig. If a gene has no previous neighbor the *s_g_*_−1_ score is 0. *f* (*g*) is a function whose result depends on the gene partition and on whether or not the gene belongs to a multigenic gene family:

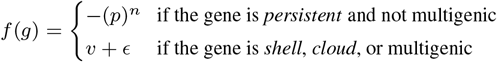

*n* is the number of consecutive *persistent* genes previously encountered and any *shell* or *cloud* gene resets its value to 0. The *p* constant is used to penalize the inclusion of *persistent* genes in an RGP whereas the *v* constant promotes the inclusion of *shell* and *cloud* genes. *E* is a constant set to 1*/*∞ that is used to obtain identical results regardless of the reading direction of the contig. A gene family is considered as multigenic if it contains duplicated genes in more that *d*% of the genomes in the pangenome graph. The default values of *p*, *v* and *d* were determined empirically and set to 3, 1 and 5%, respectively. In the case of circular sequences, if at least one gene had a score of 0, the algorithm continues after the end of the sequence to reevaluate gene scores from the beginning until reaching a gene score of 0 or reaching the last gene that had a score of 0 at the first pass.

#### 2.1.2 Detection of RGPs and score updates

After all genes have been associated to a score *s_g_*, RGPs are detected using the following algorithm on each contig.

• Step 1: initialization of a new RGP

- If no gene on the contig has a *s_g_* ≥ *s_min_*, stop here.
- Select the gene *g* with highest score *s_g_* (in case of equality, the gene closest to the end of the contig is selected).
- A new RGP that ends at the selected gene *g* is created and the score *s_g_* is assigned to the RGP.
• Step 2: extraction of the RGP

- Add previous genes to the RGP until reaching a gene with *s_g_* = 0.
- Set all the *s_g_* of the RGP genes to 0.
- Save the RGP if its length in nucleotides is ≥ *l_min_*
• Step 3: score updates

- Recompute *s_g_* scores from the gene selected at step 1 until reaching a gene with *s_g_* = 0.
- Go to step 1.

This algorithm results in the prediction of RGPs that correspond to ordered sets of genes. A minimal gene score criterion *s_min_* is used as a threshold to consider an RGP. Its default value is set to 4. In addition to *s_min_*, a minimal length in nucleotides of an RGP *l_min_* is defined and set to 3 000 by default.

#### 2.1.3 Grouping RGPs into spots

From a set of genomes with predicted RGPs, spots are determined by comparing their *persistent* flanking genes. At both ends of each RGP, we select the *c* consecutive genes that are *persistent* and not multigenic. These genes are ordered according to their distance from the RGP and then converted into their corresponding gene family. The borders of an RGP is thus defined as a pair of ordered sets of gene families. A graph *G*(*V, E*) is built where each node *v* represents the borders of an RGP and each edge indicates that they share similar sets of gene families. Two borders *v_i_* and *v_j_* are similar if their first *e* gene families are identical or if their ordered sets overlap by at least *o* families. If all borders of two compared RGPs match, we add an edge between their corresponding nodes. Once the graph is built, all connected components are extracted and corresponds to the spots. Then, the list of associated RGPs are retrieved for each spot. The default values of *c*, *e* and *o* are 3, 1 and 2, respectively. Spots are associated to multiple metrics such as the numbers of RGPs, gene families and different sets of gene families that compose the RGPs. RGPs that do not have *c* consecutive *persistent* genes on both ends are not considered for spot prediction. Either they are not complete as they end at contig borders or they are plasmids and thus do not have a *persistent* context.

### 2.2 Benchmark protocol

To assess the reliability of GI detection by panRGP, we used two previously published reference datasets (Langille *et al.*, 2008) that were recently updated (Bertelli *et al.*, 2018). The C-dataset is made of 104 genomes among 53 species from which GIs have been automatically predicted using a comparative genomic method based on IslandPick (Langille *et al.*, 2008). The L-dataset contains 6 genomes whose GIs have been curated. While it does not cover as much microbial diversity, it contains literature-curated GIs rather than automatically detected ones which makes it a much more reliable source of information. Genomes of the L-dataset are also present in the C-dataset but it should be noted that the GIs of C-dataset only partly cover those of the L-dataset (Bertelli *et al.*, 2018). For each dataset and in each genome, ‘positive regions’ correspond to regions that are potential GIs and ‘negative regions’ to those that are not considered to be GIs.

A prerequisite for the execution of panRGP is the computation of pangenomes for each studied species. All available NCBI RefSeq genomes (downloaded the 21st November 2019) (Haft *et al.*, 2017) were used as PPanGGOLiN input to obtain a partionned pangenome graph. However, only species with at least 15 RefSeq genomes could be analyzed due to the statistical constraint of the PPanGGOLiN method. Among the 54 species present in the GI datasets, 14 of them did not meet this condition. Furthermore, *Prochlorococcus marinus* was not considered in the analysis as this group of organisms does not seem to be a relevant single species at the genomic level (Parks *et al.*, 2018). An additional 3 had to be removed as their assemblies did not match the version which were originally used to predict their GIs. A total of 81 genomes from 36 species from the reference datasets could be analyzed. The list of RefSeq genomes with their identifiers that were used to build pangenomes is available from https://github.com/axbazin/panrgp_supdata. The panRGP results were obtained with the PPanGGOLiN software version 1.1.72 with default parameters.

The results of panRGP were compared to other tools on the same reference datasets. We included tools that were found to correctly predict GIs in a recent study (Bertelli *et al.*, 2018): Islandpath-dimob (Bertelli and Brinkman, 2018), SigiCFR (Waack *et al.*, 2006), SigiHMM (Waack *et al.*, 2006), Alien Hunter (Vernikos and Parkhill, 2006), PredictBias (Pundhir *et al.*, 2008), ZislandExplorer (Wei *et al.*, 2017) and IslandViewer4 (Bertelli *et al.*, 2017), which combines results from Islander (Hudson *et al.*, 2014), Islandpath-dimob, SigiHMM and IslandPick (Langille *et al.*, 2008). In addition, three recent tools for GI detection were included in the benchmark: GI-cluster (Lu and Leong, 2018), XenoGI (Bush *et al.*, 2018) and IslandCafe (Jani and Azad, 2019). GI-Cluster relies on sequence composition and functional annotation to cluster regions. XenoGI uses bidirectional best hits with the information of a phylogenetic tree to identify gene families that arose from the same HGT. IslandCafe uses sequence composition and functional annotation with a custom database of hidden Markov models including genes that are often associated with HGT. All these tools were run with default parameters. XenoGI was used only with the L-dataset as it requires a manual selection of related genomes with a phylogeny. We selected four or five genomes from closely related species and compared them with mashtree (Katz *et al.*, 2019) to obtain the phylogenetic tree for each analyzed species. The selected genomes and the trees that were used for XenoGI are available from https://github.com/axbazin/panrgp_supdata. For Islandviewer and PredictBias, the predicted GIs were downloaded from their respective websites. Sofware versions, or commit numbers, and the mode of installation are provided in https://github.com/axbazin/panrgp_supdata.

To evaluate these tools, we compared their predictions with the positive and negative regions of the two datasets using the protocol described in (Bertelli *et al.*, 2018). The predicted regions that correspond to positive regions are considered as true positives (*T P*) and those that correspond to negative regions are considered as false positives (*FP*). The negative regions that were not predicted are true negatives (*T N*) and the positive regions that were not predicted are false negatives (*FN*). We computed Matthew’s correlation coefficient (*MCC*), *F* 1*score*, *accuracy*, *precision* and *recall* as in (Bertelli *et al.*, 2018) to compare the prediction performance of the different tools.

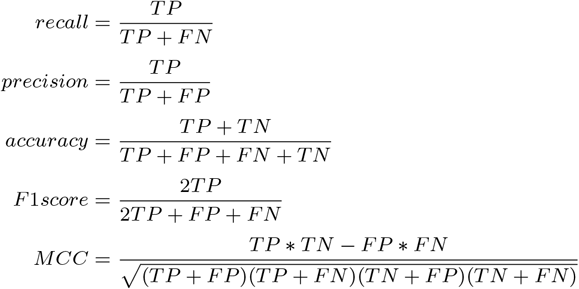

### 2.3 Preparation of the MAG dataset

In addition to the benchmark, 1416 *Escherichia coli* MAGs that have been published in a recent metagenomic study (Pasolli *et al.*, 2019) were downloaded from https://opendata.lifebit.ai/table/SGB. These MAGs were annotated using Prokka 1.13 (Seemann, 2014) before being analyzed by panRGP to predict RGPs and spots. The analysis was made with version 1.1.72 of PPanGGOLiN using the –defrag parameter to link potential gene fragments to their native gene family.

## 3 Results and Discussion

### 3.1 Software overview

The methods of panRGP for the detection of RGPs and spots have been implemented in the PPanGGOLiN pangenomic software suite (version ≥ 1.1.72) available through Github (https://github.com/labgem/PPanGGOLiN) under the CeCiLL 2.1 open source license. It is coded in Python3 with embedded code in C and can be easily installed using bioconda (Grüning *et al.*, 2018). We have also written an extensive documentation of every possible output files and the different ways of using the software in the GitHub wiki (https://github.com/labgem/PPanGGOLiN/wiki).

An overview of the panRGP workflow is given in Figure 2. The whole workflow can be run through the command ‘ppanggolin panrgp’. First, the PPanGGOLiN partitioned pangenome graph is built from a set of input genomes chosen by the user. The genomes are expected to be from the same species and can be provided as gff3/gbff annotation files or as fasta sequences. In the latter case, genomes are annotated using the procedure described in (Gautreau *et al.*, 2020). A clustering step using MMseqs2 (Steinegger and Söding, 2017) is then executed if gene families are not provided as input by the user. In that case, families are built with 80% amino acid identity and 80% coverage on both query and target proteins for the sequence alignments and with the greedy set cover algorithm for the clustering (default PPanGGOLiN parameters). Afterwards, the graph is constructed and partitioned in *persistent*, *shell* and *cloud* families. Finally, RGPs and spots are predicted using the methods described above. The pangenome and all analysis results are stored in an HDF5 file. Each step of the workflow has a dedicated subcommand in the software suite. The user can then adapt parameters or (re-)execute only a part of the workflow. These subcommands take as input either the raw original files or an HDF5 file representing the pangenome.

**Fig. 2.**
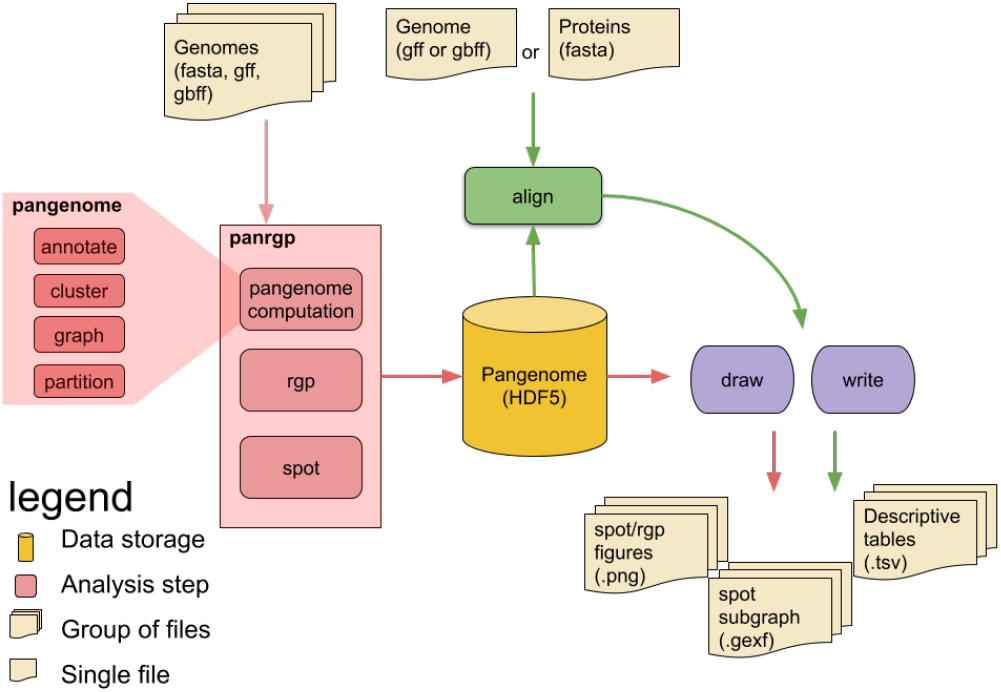
Overview of the panRGP workflow. Each rounded box represents one of the possible commands of the software. The panRGP workflow computes the pangenome using the PPanGGOLiN method, predicts the RGPs and gathers them into spots. The pangenome and results are saved in an HDF5 file. This file can be used as an input with the ‘align’ command (i) to compare a new genome to the pangenome and predict the RGPs (ii) to align protein sequences to the pangenome families to extract related RGPs and spots. A summary of results in tsv text files and figures can be obtained using respectively ‘write’ and ‘draw’ commands, respectively.

When a reference pangenome is available, a new genome of the same species can be used as input to predict RGPs using the algorithm described in section 2.1. This is done by the ‘align’ subcommand that maps the genes of the genomes to the gene families of the pangenome using the MMseqs2 software. The ‘align’ subcommand can also be used with protein sequences as input. In this case, they will be aligned to the pangenome gene families and the related RGPs and spots will be extracted, thus providing contextual information for proteins of interest.

The panRGP workflow ends by providing different output files. Tab-separated values (tsv) files contain a summary of the predicted RGPs and spots. Figures can be drawn to represent all RGPs of a spot using the genoplotR library (Guy *et al.*, 2010). Furthermore, a subgraph of the pangenome graph (in a GEXF format) can be extracted to represent all the gene families found in a spot with their genomic organization (see subsection 3.3). It can then be visualized with the Gephi software (Bastian *et al.*, 2009) by applying a layout algorithm such as ForceAtlas2 (Jacomy *et al.*, 2014).

### 3.2 Benchmark results

To evaluate the panRGP method, we ran a benchmark as described in ‘Materials and Methods’ section in comparison with ten other tools for GI prediction and on two different reference datasets. Those methods can be classified in two types: comparative-based and composition-based methods. IslandViewer4 is an hybrid method as it combines both approaches. It aggregates results from SigiHMM which is a composition-based method, and IslandPick which is a comparative-based method. Other methods like IslandCafe and GI-Cluster use additional functional information.

The C-dataset contains GIs on 81 genomes that were automatically predicted using a comparative genomic method. Results for the different methods are presented in Table 1. The panRGP method gives the best results in terms of *MCC*, *accuracy* and *precision* whereas IslandViewer4 produces better results regarding *F* 1*score* and *recall*. Overall, computed metrics for IslandViewer4 and panRGP are very close. Methods based on sequence composition do not perform as well. Some have a good *precision* (e.g. SigiCFR, IslandPath-DIMOB) but their *recall* and *accuracy* values are lower than comparative genomics based methods. It should be noted that the C-dataset was generated using comparative genomics which may explain the good performance of IslandViewer4 and panRGP, which are the only tools using comparative genomics.

**Table 1.** Benchmark results on the C-dataset

| Tool | MCC | F1 score | Accuracy | Precision | Recall | Approach |
| --- | --- | --- | --- | --- | --- | --- |
| panRGP | **0.778** | 0.809 | **0.924** | **0.949** | 0.764 | Comparative |
| IslandViewer4 | 0.762 | **0.820** | 0.911 | 0.908 | **0.788** | Hybrid |
| IslandPath-DIMOB | 0.523 | 0.570 | 0.781 | 0.891 | 0.477 | Composition |
| GI-Cluster | 0.483 | 0.592 | 0.780 | 0.674 | 0.633 | Composition / Functional |
| SigiCRF | 0.450 | 0.492 | 0.803 | 0.909 | 0.398 | Composition |
| PredictBias | 0.393 | 0.528 | 0.739 | 0.593 | 0.633 | Composition |
| IslandCafe | 0.377 | 0.444 | 0.761 | 0.769 | 0.355 | Composition / Functional |
| AlienHunter | 0.364 | 0.526 | 0.754 | 0.586 | 0.551 | Composition |
| SigiHMM | 0.338 | 0.455 | 0.756 | 0.655 | 0.376 | Composition |
| ZislandExplorer | 0.194 | 0.260 | 0.705 | 0.690 | 0.201 | Composition |

The L-dataset contains curated GIs on 6 genomes that were expertized. Results for the different methods are presented in Table 2. The benchmark was carried out on the same methods plus XenoGI which is a comparative-based method that uses a phylogenetic tree to detect insertion events from HGT. As with the C-dataset, panRGP performs best in terms of *MCC*, *accuracy* and *precision*, as well as the *F* 1*score*. Regarding the *recall*, XenoGI provides the best results. IslandViewer4 performs well but the metrics drop somewhat in comparison to the C-dataset. IslandCafe and GI-cluster which both rely on composition and functional annotation fare much better on this dataset. Overall, composition-based methods perform worse than comparative-based ones.

**Table 2.** Benchmark results on the L-dataset

| Tool | MCC | F1 score | Accuracy | Precision | Recall | Approach |
| --- | --- | --- | --- | --- | --- | --- |
| panRGP | **0.879** | **0.932** | **0.931** | **1.000** | 0.884 | Comparative |
| XenoGI | 0.829 | 0.917 | 0.905 | 0.935 | **0.924** | Comparative |
| IslandViewer4 | 0.684 | 0.791 | 0.817 | 0.998 | 0.669 | Hybrid |
| IslandCafe | 0.606 | 0.715 | 0.752 | **1.000** | 0.574 | Composition / Functional |
| GI-Cluster | 0.589 | 0.743 | 0.761 | 0.870 | 0.714 | Composition / Functional |
| PredictBias | 0.587 | 0.805 | 0.788 | 0.856 | 0.771 | Composition |
| IslandPath-DIMOB | 0.527 | 0.636 | 0.702 | 0.998 | 0.479 | Composition |
| SigiCRF | 0.424 | 0.520 | 0.687 | 0.993 | 0.434 | Composition |
| AlienHunter | 0.398 | 0.642 | 0.705 | 0.753 | 0.570 | Composition |
| SigiHMM | 0.268 | 0.444 | 0.591 | 0.817 | 0.325 | Composition |
| ZislandExplorer | 0.163 | 0.278 | 0.513 | 0.833 | 0.180 | Composition |

This benchmark shows that comparative-based methods are the most reliable to predict GIs in comparison to composition-based methods. Surprisingly, IslandViewer4 that combines both approaches does not perform better than panRGP or XenoGI, which only rely on comparative genomics. We can assume that the tools based on comparative genomics are more reliable when they use a larger number of genomes representing a greater diversity within the studied species. This may explain why panRGP comes on top in this study. Indeed, the reference pangenome graph that is used by panRGP can contain information from hundreds to thousands of genomes. On the other hand, IslandViewer4 and XenoGI are limited to few genomes. IslandViewer4 uses up to 12 genomes (6 by default). For XenoGI, authors indicate the use of 500 gigabytes (GB) of memory and a run time of 20 hours on 50 threads for 40 strains, thus limiting its use on larger datasets. Conversely, panRGP can use far more than hundreds of genomes without requiring extensive computational resources. For the complete workflow of panRGP including the pangenome construction and RGP/spot prediction, it takes 3 minutes and 1.2 GB of memory to analyze 40 strains of *E. coli* and 45 minutes and 14 GB of memory for 1000 strains of *E. coli* on 16 threads of an Intel Xeon CPU E5-2699v3. Most of the time is dedicated to genome annotation and pangenome partitioning.

### 3.3 Application of panRGP on a pangenome built from MAGs

To illustrate the potential of panRGP on MAGs, we studied the genomic context of a previously described hotspot in *E. coli* (Lescat *et al.*, 2009) using a pangenome constructed from MAG sequences from a recently published metagenome dataset (Pasolli *et al.*, 2019). The pangenome was built using 1 413 MAGs. It is made of 43 741 gene families including 5 111 342 genes. Those families are partitioned into 3 724 *persistent*, 2 490 *shell* and 37 618 *cloud* gene families. The *persistent* genome is very similar to the one that was already computed on a pangenome built from GenBank genomes (3706 families) (Gautreau *et al.*, 2020) indicating that the *persistent* was well retrieved even though the MAGs are much more fragmented and incomplete than genomes from isolates. As a consequence, pangenome partitions obtained on these MAGs can be a reliable source of information to predict GIs with panRGP. A total of 47 692 regions were predicted by panRGP among which 18 030 are in-between *persistent* genes and could be used to predict spots of insertion with the algorithm previously described. Those 18 030 RGPs have been grouped into 294 spots of insertion. A list of all spots with descriptive metrics are available from https://github.com/axbazin/panrgp_supdata..

Among all the predicted spots, we focused our analysis on the *leuX* tRNA hotspot as it is one of the most diverse region of *E. coli* and involved in pathogenicity (Blum *et al.*, 1994; Touchon *et al.*, 2009). This spot was described as being in-between two core genes, namely *uxuA* and *ahr* (previously named *yjgB*), in a comparative analysis of 14 genomes (Lescat *et al.*, 2009). To retrieve this spot from panRGP results, we used the protein sequences of both of those genes from *E. coli* K-12 MG1655 (P24215 and P27250 proteins from UniProt, (UniProt Consortium, 2019)) and aligned them to the pangenome gene families (subcommand ‘align’ see ‘Materials and Methods’ section). We find that only one panRGP spot is associated with both genes and corresponds to the spot number 10. It gathers 131 RGPs that are represented by 79 different sets of gene families. The size of these RGPs is between 5 to 91 genes, with an average of 19 genes. There is a total of 585 different gene families. Among all predicted spots, it is the third most diverse in gene family content, confirming that it is one of the most dynamic regions of the *E. coli* genome.

Figure 3 shows the pangenome subgraph of this spot in a compact representation with the indication of gene names that are most often associated with each family. The spot borders were formerly assumed to be *ahr* and *uxuA* genes (Lescat *et al.*, 2009). While we agree that *ahr* borders the spot, the *nan* operon composed of the *nanS*, *nanM* and *nanC* genes (previously named *yjhS*, *yjhT* and *yjhA*) is the most common border predicted by panRGP instead of *uxuA*. Indeed, *nan* genes are *persistent* and thus present in most *E. coli* genomes from the gut microbiome. The punctual deletion of the fimbrial operon *fim* in few *E. coli* strains (e.g. 55989 and O42 strains, see Figure 4 in (Lescat *et al.*, 2009)), which is located between the *nan* operon and *uxuA* in the other strains, misled the authors in determining the hotspot frontiers as few genomes where available when their work was published.

**Fig. 3.**
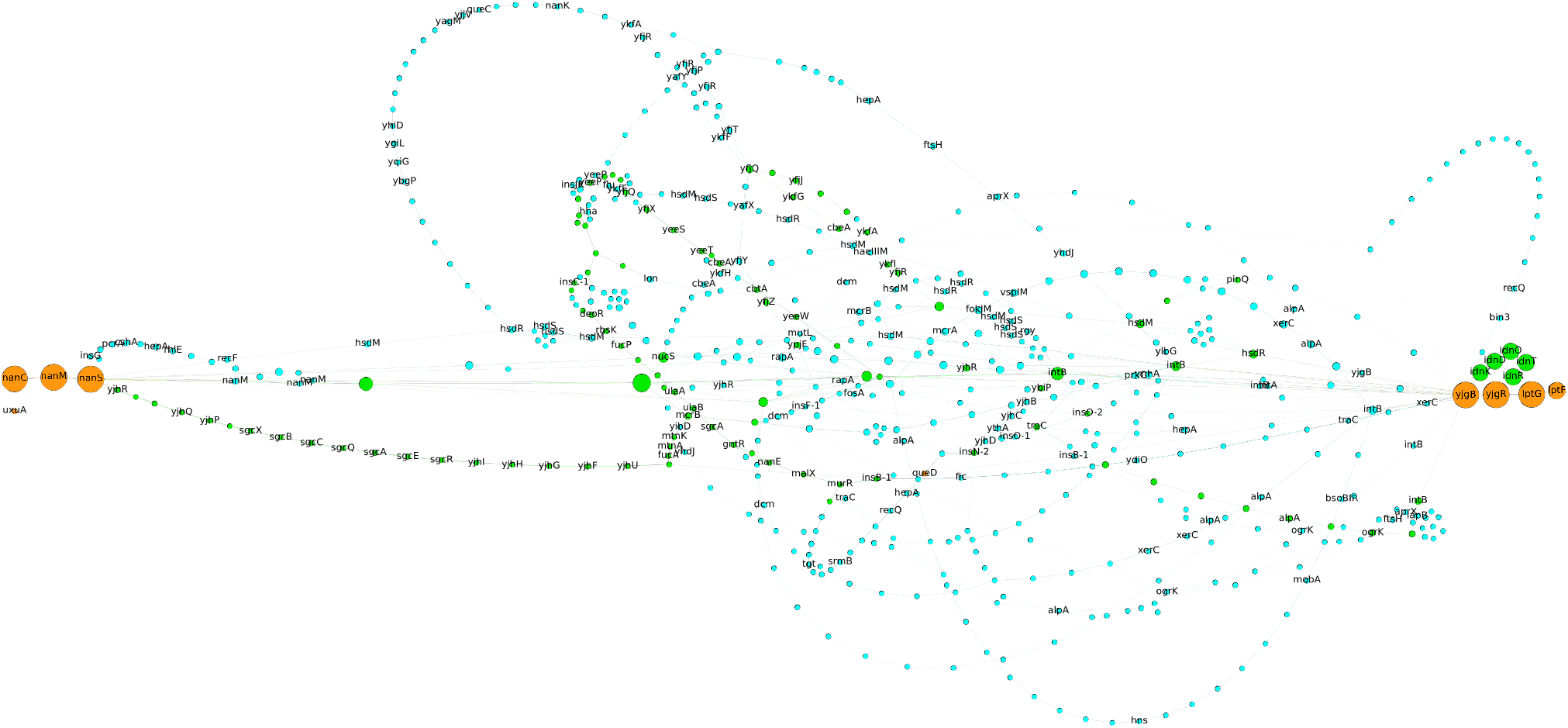
Pangenome subgraph of the *leuX* hotspot in MAGs of *E. coli*. This figure illustrates the genetic diversity of the leuX hotspot both in terms of gene content and genome organization. Each node is a gene family and an edge is drawn between two families that share neighboring genes. Colors for persistent, shell and cloud pangenomes partitions are orange, green and blue, respectively. The size of the nodes is proportional to the number of occurrences in the spot. For each family, the most represented gene name is indicated. The tRNA leuX and the fim operon are not represented as they are not members of spot predicted by panRGP. leuX is located next to yjgB and the fim operon is just before nanC. This visualization was produced using Gephi (Bastian et al., 2009) and the ForceAtlas2 layout algorithm (Jacomy et al., 2014).

The spot detection method illustrated here is inspired from (Oliveira *et al.*, 2017) with, however, a fundamental difference: our method is not centered on a pivot genome but is applied on the whole pangenome without any reference. Furthermore, it allows for variations in terms of gene content and organization in the definition of the spot borders. The *leuX* hotspot is a great example showing that spot borders can vary throughout the evolution of a species. The panRGP method provides an exhaustive list of spot associated with several metrics (e.g. numbers of RGPs, gene families and different sets of families). Those results can be the starting point of studies on the dynamics of GIs within and between species.

## 4 Conclusion

We presented an original method that can identify RGPs on thousands of genomes and analyze them together to detect spots of insertion. Indeed, panRGP uses a partitioned pangenome graph of gene families that makes comparative-based approach to predict GIs more efficient. Indeed, our method is much more scalable on large datasets than already published tools, which rely on time-consuming pairwise sequence comparisons. We showed that panRGP results are highly reliable when compared to a dataset of curated GIs. We introduced a novel algorithm for the detection of spots of insertion which was illustrated in the context of the analysis of an *E. coli* hotspot using MAGs from the human gut. Overall we believe that panRGP provides an original approach to detect GIs to study their diversity and dynamics in a species of interest. Its ability to predict GIs and spots among thousands of genomes makes it an ideal approach for large scale studies.

The tool is freely available and easily installable as part of the PPanGGOLiN software suite. It is also integrated in the MicroScope platform with a dedicated web page for result analysis and exploration of prokaryotic genomes (Vallenet *et al.*, 2019). An improvement of panRGP could be to analyze conserved alternative paths within RGPs using the pangenome graph structure. This could allow to automatically identify functional modules, i.e. set of genes involved in the same biological process akin to what was described in (Lescat *et al.*, 2009) and (Touchon *et al.*, 2009).

## Acknowledgements

Valentin Sabatet for initial work and proof of concept on studying plastic regions using pangenome partitions. Mark Stam and Mathieu Dubois for insightful discussions. Mylène Beuvin for her artistic sense.

